# Task sharing clubfoot treatment in Latin America: a cross-sectional survey of expert opinions

**DOI:** 10.1101/755298

**Authors:** Roxana Martinez, Tracey Smythe, Christie Pettitt-Schieber, Jennifer Everhart, Joshua Hyman

## Abstract

While the Ponseti method has quickly become the mainstay of clubfoot treatment in most parts of the world, its dissemination and successful implementation in Latin America has been more limited. The additional shortage of orthopedic surgeons in this region makes task sharing a practical approach to address gaps in service provision. We designed an online survey to assess needs, perceptions, and willingness to task share the delivery of the Ponseti method by Ponseti-method-trained physicians across Latin America. Multiple-response questions were summarized and an applied thematic analysis approach was used to analyze free-response questions. We achieved a 66% response rate (31 of 47 experts responded). Our findings illustrate that most physicians feel the need for disseminating and improving Ponseti training, as well as having additional support for clubfoot treatment. While physicians who treat clubfoot have mixed opinions on the role of nonphysicians treating clubfoot, most report logistical concerns and insufficient training as barriers to their inclusion. Given this and the need for improved, more accessible clubfoot care across Latin America, future clubfoot treatment efforts may benefit from incorporating task sharing between orthopedic surgeons and non-physician personnel.

## Introduction

Clubfoot, defined as the downward- and inward-turning of the foot, is one of the most common musculoskeletal birth deformities in the world (1). When left untreated, clubfoot may cause lifelong physical impairment, social isolation, and economic deprivation. The global paradigm for management of clubfoot has shifted from the provision of extensive surgical correction to implementation of a minimally-invasive, conservative correction that is both low-cost and highly-effective; this technique is called the Ponseti method (2).

The Ponseti method, created by Dr. Ignacio Ponseti, involves a correction and a maintenance phase (3). The correction phase comprises serial manipulations with simultaneous correction of the four components of the deformity: cavus, adductus, varus, and equinus. A series of long-leg plaster of Paris casts hold the corrected foot position, usually followed by an outpatient Achilles tenotomy. Immediately after the removal of the final cast, the corrected foot is placed in a foot abduction brace (FAB) with the aim of preventing recurrence. The FAB should be worn 23 hours a day for the first three months and then only at night until the age of four years; the FAB itself is changed as the child’s foot grows. Both the maintenance and correction phases are equally important for success of clubfoot management. Although success rates of the Ponseti method vary based on when treatment is initiated, patient adherence, and provider experience, complete correction can be achieved in the majority of patients with success rates as high as 95% (4).

While the Ponseti method has quickly become the mainstay of clubfoot treatment in most parts of the world – US, UK, Australia, India, and parts of Africa – its dissemination and successful implementation in Latin America has been slower (4). Despite clubfoot being reported as one of the most commonly encountered pediatric orthopedic conditions in parts of Latin America, when compared with patients in higher income nations, patients have a later age of presentation, more time spent in the manipulation and casting phase, lower rates of tenotomy, and higher rates of relapse (5, 6). Inefficient healthcare systems may be partly responsible for this inequity and shortage of trained orthopedic surgeons additionally contributes to barriers to care (4, 7–9). Similar to other specialty services, most orthopedic professionals work in urban population centers, further disadvantaging patients in rural areas.

Although training more orthopedic surgeons appears to be the most straightforward solution, this task requires intensive resources, time, and effort, making it an unrealistic short-term plan (10). Instead, more radical reform of specialist services is needed, particularly where decentralization or integration of services into primary care is necessary to improve access to care and achieve universal health coverage. Task sharing is one practical approach to addressing such gaps in human resources; it involves teaching competencies previously held by specialists to other personnel (11, 12). Originally created by the Lancet Commission on Global Surgery (LCoGS), task sharing –a practice whereby nonsurgeon professionals and clinicians are trained to do simple procedures, whilst having access to surgical professionals who normally do said procedures – is commonly used both in high- and low-resource settings, having been found to be safe and cost-effective (10). In clubfoot treatment, findings from Malawi illustrate that non-physician staff who received training in the Ponseti method were at least as effective as physicians (13). Similar findings were found in Nepal, Vietnam, United States, United Kingdom, and Canada (13–18). Moreover, treatment by physiotherapists in the United Kingdom provided lower rates of additional treatment (14% vs. 26%, p=0.075) as well as a lower rate of additional procedures (6% vs. 18%, p=0.025) when compared to physician-directed groups (17).

Despite the promising data available on the use of nonphysician personnel to implement the Ponseti technique and the shortage of orthopedic surgeons in Latin America, no literature exists on task sharing the Ponseti method for the Latin America region. Given the cost-effectiveness of the Ponseti method, the barriers identified by physicians and caregivers, and the shortage of orthopedic surgeons in Latin America (4, 7, 8), this study aims to answer two questions: (1) Do non-physician personnel (e.g. physical therapists, nurses, etc.) have a role in treating clubfoot, such as assisting with manipulation and casting, providing education, or other tasks? and (2) what are the perceptions and attitudes from Ponseti-trained physicians regarding nonphysician providers implementing (i.e. task sharing) the Ponseti method?

## Materials and methods

### Ethics

This study was determined to be exempt from Interview Review Board approval by the Columbia University Administrative Review Committee (IRB-AAAS3714).

### Study and questionnaire

The survey in this study was designed to assess needs, perceptions, and willingness to task share the Ponseti method by Ponseti-trained physicians across Latin America. It consisted of 17 questions, including multiple response and free-response questions (see Appendix 1). All questions were reviewed by the contributing authors. Questions were translated to Spanish by a native speaker and read by other native Spanish speakers for clarity. Questions were uploaded to Qualtrics, an online survey platform.

Basic demographics such as occupation, country of employment, age, and gender were recorded. Questions to gauge experience with treating clubfoot were asked, such as how participants were trained, length of time treating clubfoot, and their monthly estimate of new clubfoot patients in their clinic. Adherence to and success rates of the Ponseti method were evaluated for each provider through questions about percentage of clubfoot patients completing casting and/or bracing and rate of patients receiving a tenotomy. Participants were asked in a multiple response question to identify barriers to implementing the Ponseti method, and these barriers were divided into ten categories (Table 2). Participants were asked to rank, on a five-point Likert scale, the extent to which they agreed with several statements regarding time, resources, and attitude towards teaching patients about clubfoot. Participants were additionally asked to identify collaborators for implementing the Ponseti method (Table 3), and were given the opportunity to elaborate in a free-response question on nonphysician staff collaboration with physicians. They were asked to share their opinion of nonphysician staff having a role in treating clubfoot, and then asked to elaborate in a free-response question. A final, open-ended question asking for further suggestions for improving clubfoot treatment was included.

### Participants and study procedures

A list of physicians with known experience in clubfoot treatment was obtained from an international non-profit organization. Participants were contacted via email a total of three times at one-week intervals.

Surveys were distributed to 47 practitioners using the Qualtrics platform from April 2019 to June 2019. All survey responses were anonymous.

### Statistical analyses

Multiple-response questions were summarized and graphed using Qualtrics. An applied thematic analysis approach was used to draw results from free-response questions. An initial code structure was devised for each free-response question and applied systematically to each free-text response.

## Results

### Demographics

Thirty-one practitioners responded to the online survey, yielding a response rate of 66% and including 29 orthopedic surgeons, one pediatrician, and one unspecified physician. Participants were recruited from eight countries as outlined in Table 1. The average age of respondent was 40 years, with 71% of respondents being male. All participants had been trained in the Ponseti method. Most had been formally trained in residency (n=19, 38%) and/or a formal workshop (n=18, 36%). Some had one-on-one training from orthopedic surgeons (n=5, 10%) and two (6%) were exclusively trained in this manner, one by Dr. Ponseti himself. Four physicians (8%) supplemented their training with online resources, while only one participant (3.2%) exclusively used online resources for training.

**Table 1.**
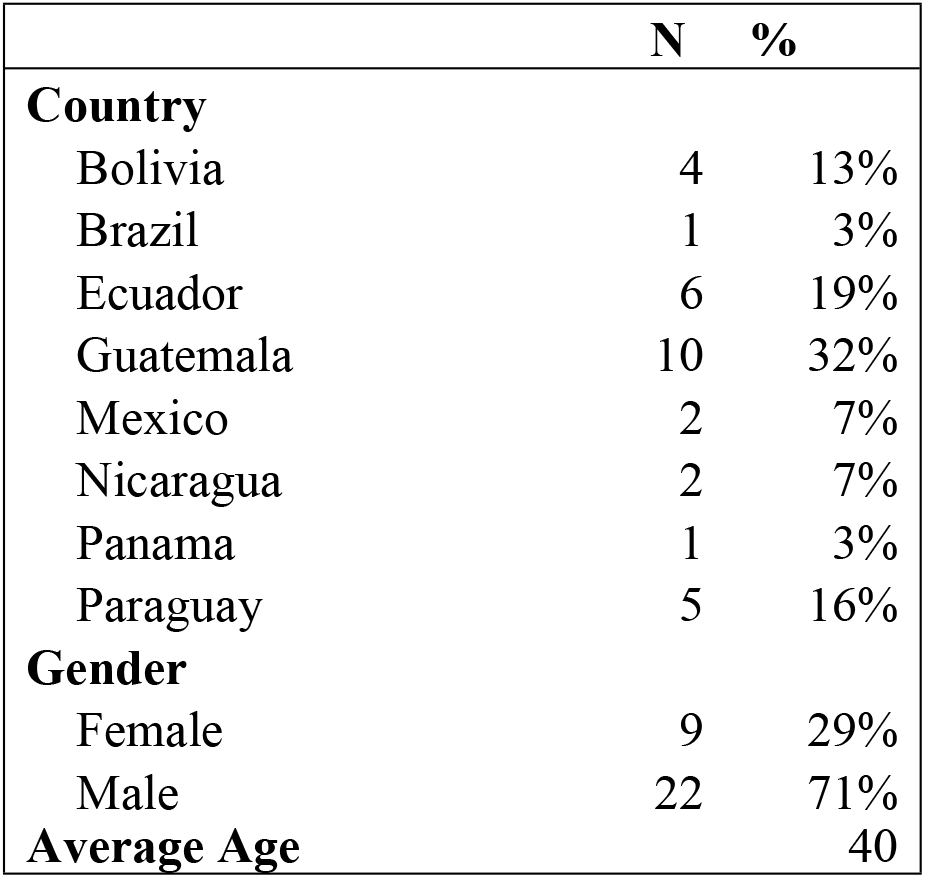
Demographics.

### Experience with treating clubfoot and clubfoot practices

The majority of physicians reported treating clubfoot for greater than five years (n=16, 51.6%), with several physicians (n=10, 32.3%) with 2-5 years of experience and some (n=5, 16.1%) with < 2 years of experience. Over half of the physicians (n=17, 54.8%) reported treating between 0 and 3 new clubfoot patients a month, while several (n=12, 38.7%) reported between 4 and 10 new clubfoot patients a month and only two (6.5%) reported more than 10 a month. The majority of physicians (n=21, 67.7%) reported that, within the past year, more than two-thirds of their patients completed the casting phase; eight (25.8%) reported a fraction between one-third and two-thirds and only two (6.5%) reported a fraction of less than one-third. Most respondents (n=17, 54.8%) reported a rate of over two-thirds for completing both casting and bracing; eight (25.8%) reported a rate between one-third and two-thirds and the remaining (n=6, 19.4%) reported a completion rate of less than one-third. The majority (n=22, 70.1%) reported that over two-thirds of patients obtained a tenotomy; seven respondents (22.6%) reported a rate between one- and two-thirds and the remainder (n=2, 6.5%) reported a rate of less than one-third.

### Barriers to implementing the Ponseti method

The majority of physicians (n=25, 80.6%) agreed or somewhat agreed with the statement, “I have all the material resources that I need to implement the Ponseti method” (Fig 1). The remaining disagreed or somewhat disagreed (n=6, 19.4%). Similarly, the majority of physicians (n=26, 83.9%) agreed or somewhat agreed that physicians have enough time to successfully implement the Ponseti method; four (12.9%) disagreed or somewhat disagreed and one (3.2%) was neutral (Fig 2). All agreed or somewhat agreed with the statement that teaching caregivers, parents, and patients is an important part of the physician’s job.

**Fig 1.**
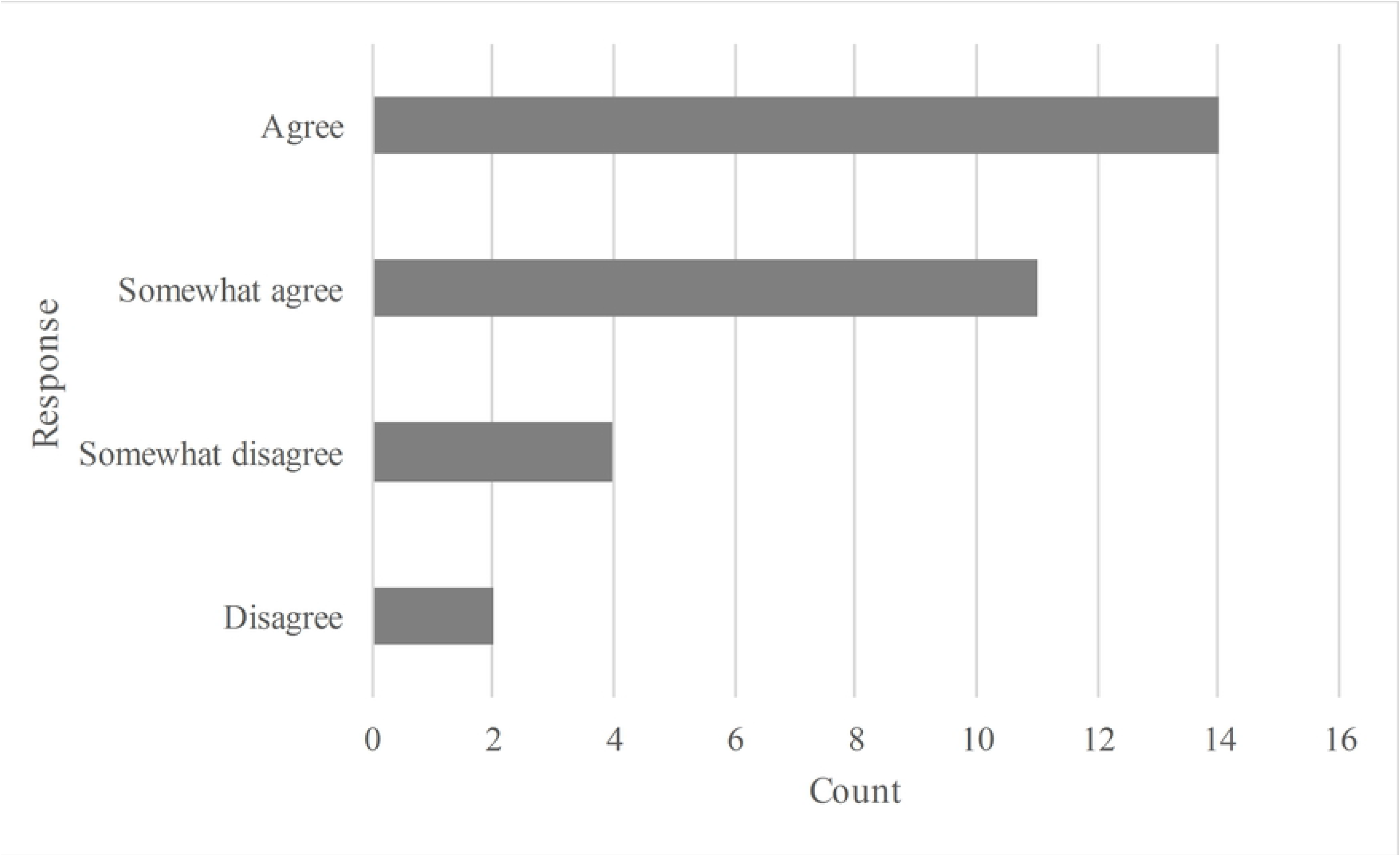
Count of responses to “I have all the material resources that I need to implement the Ponseti method.”

**Fig 2.**
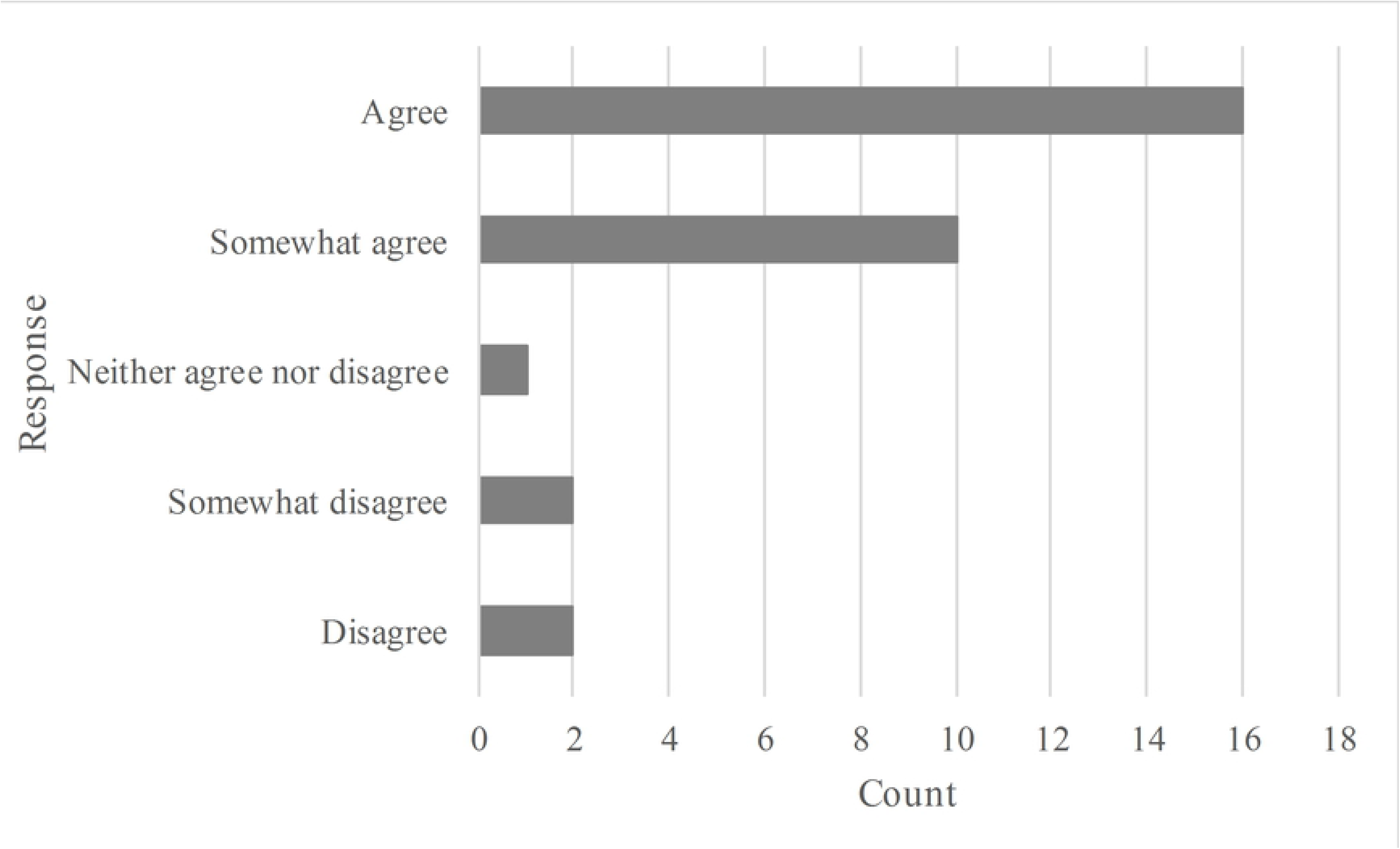
Count of responses to “I have the time necessary to successfully implement the Ponseti method, including the casting phase.”

Common barriers to clubfoot treatment are reported in Table 2. For this multiple response question, physicians identified an average of 2.8 barriers, the most common being that parents stopped using the brace, (n=23, reported by 74.2% of physicians).

**Table 2.** Reported barriers to clubfoot treatment

| Answer | Count of Physicians Who Chose Response | Percentage of Physicians Who Chose Response |
| --- | --- | --- |
| Parents stopped using brace | 23 | 74.2 |
| Failure of patients to return to clinic | 18 | 58.1 |
| Lack of or difficulty obtaining braces | 14 | 45.2 |
| Cost of treatment to the patient | 12 | 38.7 |
| Lack of or inadequate casting materials | 6 | 19.4 |
| Insufficient reimbursement to physicians | 5 | 16.1 |
| Inadequate training in Ponseti method | 5 | 16.1 |
| Inability of child to tolerate bracing | 2 | 6.5 |
| Insufficient reimbursement for cost of supplies | 2 | 6.5 |
| Insufficient reimbursement to other staff | 1 | 3.2 |
| Total Responses | 88 |  |

### Role of nonphysician staff in clubfoot treatment

Most physicians reported collaborating with others to treat clubfoot, most commonly other orthopedic surgeons, nurses, physical therapists/physiotherapist, general practitioners, and medical assistants (Table 3).

**Table 3:**
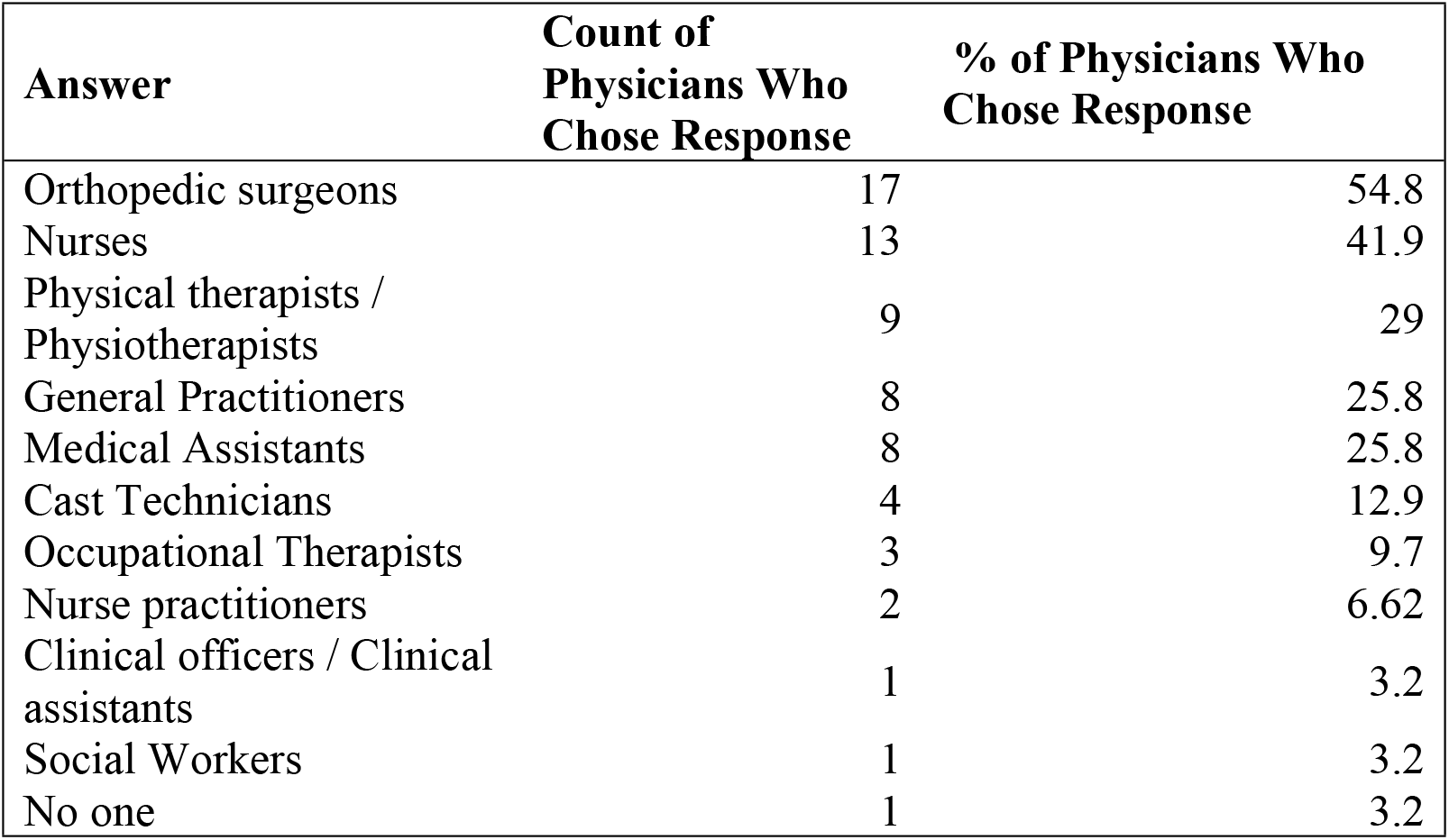
Number of physicians identifying the following staff members as collaborators for treatment

| **Answer** | **Count of Physicians Who Chose Response** | **% of Physicians Who Chose Response** |
| --- | --- | --- |
| Orthopedic surgeons | 17 | 54.8 |
| Nurses | 13 | 41.9 |
| Physical therapists / Physiotherapists | 9 | 29 |
| General Practitioners | 8 | 25.8 |
| Medical Assistants | 8 | 25.8 |
| Cast Technicians | 4 | 12.9 |
| Occupational Therapists | 3 | 9.7 |
| Nurse practitioners | 2 | 6.62 |
| Clinical officers / Clinical assistants | 1 | 3.2 |
| Social Workers | 1 | 3.2 |
| No one | 1 | 3.2 |

Free text responses to how assistants collaborate with surveyed physicians had several recurring, major themes. The most prevalent theme (n=17, 54.8%) was that staff members assist with cast application and/or cast removal. One physician added, “I work with a cast technician and nurse. They help me with the treatment…I show them the [Ponseti] method” [Participant 24], illustrating an apprenticeship-like model between the physician and their assistants. The other prominent theme (n=13, 41.9%) that arose is that educating caregivers is fundamental. One participant stressed, “The education of parents about the regimen of the plaster and the bracing stage is extremely important” [Participant 26]. Only two (6.5%) participants commented on the importance of non-physician staff assisting by obtaining casting and bracing supplies.

Fourteen surveyed physicians (45%) believe there is a role for nonphysician staff in clubfoot treatment, while 13 (42%) did not believe there was an appropriate role, and four (12.9%) were unsure (Fig 3). Of physicians who do believe nonphysician staff have a role, the most prominent theme (n=10, 32.3%) that emerged from free-text explanations was that there is a need for collaboration when implementing the Ponseti method:

**Fig 3.**
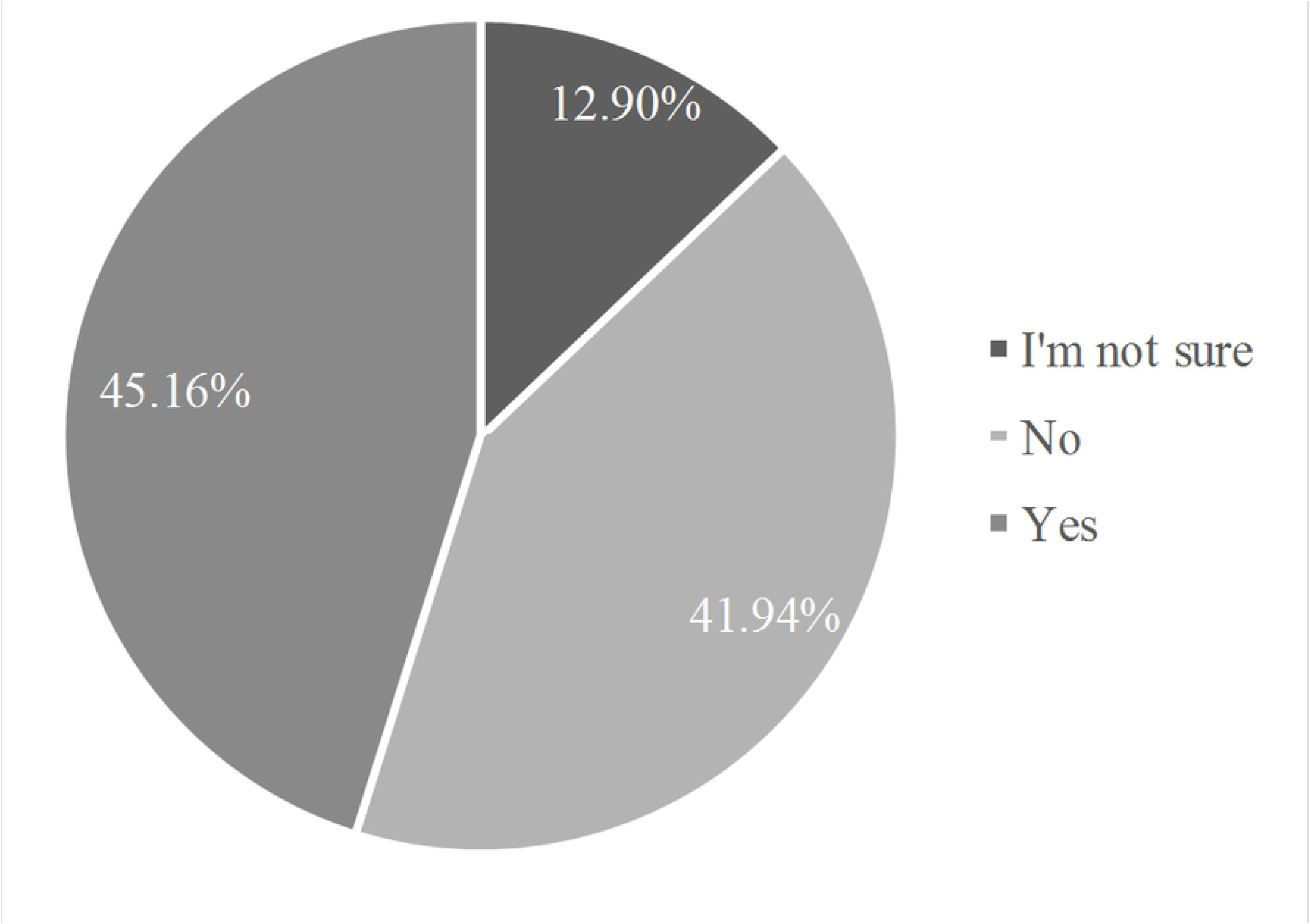
Percentage of responses to “Is there a role for paramedical staff in treating clubfoot?”

*“For the casting, two are required…” – [Participant 23]*

*“It is a teamwork. The success of a good plastering will also depend on the collaboration between plasterers.” – [Participant 19]*

Other physicians expanded on this idea, stressing the need for professionals outside of orthopedic surgery:

*“The treatment involves a lot of factors beyond simple corrections of the deformity…It requires solutions/alternatives of various areas beyond medicine.” – [Participant 20]*

*“…This treatment requires an integral approach, therefore several professionals in health are involved, as well as administrative...” – [Participant 11]*

In addition, some physicians (n=3, 9.7%) elaborated on the assistance non-physicians can provide in educating patients about clubfoot. One participant (3.2%) stated that “it’s important that each of the people who are interested in the method…those that form any part of medical careers…receive the appropriate training” [Participant 24], advocating for the dissemination of the Ponseti method for all healthcare practitioners who wish to learn. Two respondents (6.5%) commented on how nonphysician staff can assist with building rapport with patients, such as “providing assistance in doubts about the treatment or difficulties presented” [Participant 32] or “extend[ing] contact with parents and follow-up” [Participant 23].

Physicians who felt there was no role for nonphysician staff in treating clubfoot had responses that mostly focused on the theme of logistical matters (n=9, 29%). Three respondents (9.7%) explicitly stated that they were understaffed and did not have assistants, while others (n=4, 12.9%) focused more on systematic barriers:

*“The system doesn’t allow for it.” – [Participant 7]*

*“[there is] lack of coordination within the health network” – [Participant 6]*

*“…in the hospital where I work, nursing is not allowed to help in these types of procedures, even if they are needed.” – [Participant 12]*

*“I schedule the appointment of my patients.”’ – [Participant 30]*

Two participants (6.5%) focused on the challenges that arise from inconsistent placement of auxiliary personnel, stating that “there is no specific role for personnel since they take turns rotating through different services every month” [Participant 26]. Only one participant (3.2%) explicitly stated that implementing the Ponseti method should be strictly limited to doctors.

Of the physicians that were unsure of the role of nonphysician staff in treating clubfoot (n=4, 12.9%), some (n=2, 6.5%) reported uncertainty because there were no assistants available in their hospital and/or they were unfamiliar with the nonspecific essential function of nonphysician practitioners. Others (n=2, 6.5%) commented on the commitment and responsibility needed from nonphysician staff.

The most prominent theme that emerged from participants’ responses for how clubfoot treatment could be improved in their country was improving the diffusion of Ponseti training (n=18, 58.1%). In specific, some physicians called for “continuous training” [Participant 24] in addition to training being “taught in an organized manner” [Participant 30]. Some physicians (n=5, 16.1%) additionally called for changes at the healthcare system level that would assist in the diffusion of the Ponseti method by, for example:

*“…train[ing] at the primary care level to recognize clubfoot that requires the Ponseti method…creat[ing] new Ponseti Clinics and includ[ing] them in the Ministry of Health programs.” – [Participant 23]*

*“…implementing [Ponseti] in the national pediatric care program…[as] it should be part of the evaluation and comprehensive management of child’s health”-[Participant 19]*

*“…establish[ing] an efficient flow (order) of taking people or things to the treatment center…”-[Participant 20]*

*“…demonstrating the results, making them public…mak[ing] it possible to have the Ministry of Health commit to the treatment” [Participant 21]*

Similar to providing better, more systemically available training on the Ponseti method, another major theme (n=7, 22.6%) was that a greater quantity of personnel for treating clubfoot is needed. For example, one physician commented that they “have to make time to practice the Ponseti method” [Participant 27] since they are responsible for all urgent cases presenting to their hospital, illustrating that no other personnel are available to do both. Another participant stressed how “it would be ideal to be able to count on a nursing staff for consultations” [Participant 13]. These comments reiterate the need for additional staff for clubfoot care.

## Discussion

This study is the first survey to assess the role of, and orthopaedic surgeon’s attitudes towards, task sharing the Ponseti method in Latin America. Our results illustrate that physicians are equally divided on whether there is a role for nonphysician staff in clubfoot treatment, with most who are unsure or doubtful indicating primarily logistical concerns (e.g. the quantity of nonphysician assistants and the quality of their training) as barriers. However, there is consensus amongst the providers for disseminating and improving the Ponseti training in order to expand access to clubfoot treatment.

With new methods of information dissemination such as low-bandwidth training sessions on cellphones, mHealth applications, and nationwide training programs (e.g. Brazil’s standardized, national “Ponseti Brazil” workshops), clubfoot patient volume and services are likely to increase (19, 20). However, the current volume of orthopedic surgeons in Latin America trained in the Ponseti method will unlikely be able to deliver and maintain high-quality clubfoot treatment with this increasing demand for services. Our results illustrate that many physicians see at least one child with clubfoot weekly; while this is a small number, the chronicity and natural progression of clubfoot multiplies the effect that one clubfoot patient has when compared to acute, non-progressive conditions.

An additional finding from this survey is that there is low caregiver compliance with bracing and high rate of failure to return to clinic. Both findings act as major barriers to treatment and have been mirrored in previous studies (4, 7, 8). Studies from New Mexico and New Zealand, for example, have found that lack of adherence with bracing is the largest risk factor for clubfoot recurrence (21, 22). Nonphysician providers trained in the Ponseti method could potentially improve bracing adherence through, for example, home visits, for patients who need bracing care. Nonphysician providers thus could provide a supplemental and novel role in task sharing clubfoot care, and their involvement could facilitate both treatment and prevention of recurrence. Moreover, as patient education was routinely listed as one of the main barriers to successful treatment, Ponseti-trained nonphysician personnel could facilitate the spread of the Ponseti method and expand access to treatment by educating and empowering patients and their caregivers to receive optimal care (5, 7, 8). In short, sharing the task of providing caregiver education may alleviate the burden disproportionally placed on a scarce supply of orthopedic surgeons.

The results from this survey demonstrate that some physicians are amenable to having nonphysician staff assist in clubfoot treatment. We conclude that nonphysician personnel may provide human resources to fill the growing clubfoot treatment gap. While ensuring resources are not diverted away from surgical specialists, adequate support and training are needed in task sharing to ensure that (1) quality of care is maintained and that (2) the task sharing initiative can be scaled up adequately (10). Still, training alone is not enough; continued supervision is needed to ensure that non-specialist staff are confident in their tasks, carrying them out to a high quality and being able to ask questions as necessary. Adequate recognition and remuneration are additionally important to maintain motivation for non-specialist staff in carrying out their new tasks. Ultimately, task sharing – for the Ponseti method or any other clinical treatment – requires clear communication and mechanisms to support monitoring, supervision and evaluation.

### Strengths and limitations

Strengths of this study include the geographic diversity of the physicians that provided data, which may allow for broad implications of the results. Survey anonymity additionally adds to the quality and subjectivity of the data collected. Study limitations include having a small study sample size, in addition to potential selection bias given the mode of survey distribution (e.g. online, via email).

## Conclusion

In Latin America, many clubfoot treatment providers report collaborating with non-physicians to implement the Ponseti method. While physicians who treat clubfoot have mixed opinions on the role of nonphysicians treating clubfoot, most report logistical concerns and insufficient training as barriers. Given this and the need for better, more accessible clubfoot care across Latin America, future clubfoot treatment efforts may benefit from incorporating task sharing between orthopedic surgeons and non-physician personnel.

## Acknowledgements

We would like to thank MiracleFeet for their guidance and assistance in completing this project.

